# SARS-CoV-2 causes severe alveolar inflammation and barrier dysfunction

**DOI:** 10.1101/2020.08.31.276725

**Authors:** Stefanie Deinhardt-Emmer, Sarah Böttcher, Clio Häring, Liane Giebeler, Andreas Henke, Roland Zell, Franziska Hornung, Christian Brandt, Mike Marquet, Alexander S. Mosig, Mathias W. Pletz, Michael Schacke, Jürgen Rödel, Regine Heller, Sandor Nietzsche, Bettina Löffler, Christina Ehrhardt

## Abstract

Infections with SARS-CoV-2 lead to mild to severe coronavirus disease-19 (COVID-19) with systemic symptoms. Although the viral infection originates in the respiratory system, it is unclear how the virus can overcome the alveolar barrier, which is observed in severe COVID-19 disease courses.

To elucidate the viral effects on the barrier integrity and immune reactions, we used mono-cell culture systems and a complex human alveolus-on-a-chip model composed of epithelial, endothelial, and mononuclear cells.

Our data show that SARS-CoV-2 efficiently infected epithelial cells with high viral loads and inflammatory response, including the interferon expression. By contrast, the adjacent endothelial layer was no infected and did neither show productive virus replication or interferon release. With prolonged infection, both cell types are damaged, and the barrier function is deteriorated, allowing the viral particles to overbear.

In our study, we demonstrate that although SARS-CoV-2 is dependent on the epithelium for efficient replication, the neighboring endothelial cells are affected, e.g., by the epithelial cytokine release, which results in the damage of the alveolar barrier function and viral dissemination.

## INTRODUCTION

The novel severe acute respiratory syndrome coronavirus (SARS-CoV-2) is a highly pathogenic virus causing severe respiratory infections, described as coronavirus disease-19 (COVID-19) (Bar-On et al., 2020). Patients suffer from various symptoms as fever, cough, breath shortness, headache, muscle aches, and gastrointestinal symptoms. Hallmarks of severe COVID-19 courses are pneumonia, pulmonary edema, acute respiratory distress syndrome (ARDS), and multiple organ failure. In most patients, the disease has a mild course, but in some cases, e.g., elderly with comorbidities, the infection can develop into a life-threatening condition. In particular preexisting lung pathologies and systemic diseases such as diabetes predispose to severe infection courses described by George et al. (2020).

Clinical studies revealed that the virus primarily replicates in the lung, which can cause severe lung damage up to necrotic destruction of large areas of the lung tissue (Carsana et al., 2020). In autopsies of deceased COVID-19 patients, it has been observed that particularly in severe cases viral particles can disseminate throughout the body (Deinhardt-Emmer et al., 2020b; Wichmann et al., 2020). Additionally, systemic complications have been reported, such as the massive release of proinflammatory cytokines and thromboembolic events in various organs (Becker, 2020). Consequently, SARS-CoV-2 is regarded as a pneumotropic virus that infects the patient via the lung but can also cause a systemic infection that affects different organs with a high mortality rate.

Little is known about the initial infection process in the alveolar lung tissue, particularly about mechanisms that destroy the lung and mechanisms that allow the virus to affect different organs in the body. Infection models that closely reflect the patient’s situation are mainly lacking in part due to the challenges to infect mice and the difficulty of accessing and analyzing infected human alveolar lung cells. Up to now, SARS-CoV-2 infection models have been mainly performed with human airway (non-alveolar) cells or non-human cell lines that naturally express the ACE2 viral receptor, such as the African Green Monkey Vero 76 cell line (Hoffmann et al., 2020). These cells lack organ- and species-specific characteristics of human lung epithelial cells. For this purpose, cancerous lung epithelial cells (Calu-3 cells), can at least to some extent, reflect the response of the lung epithelium to viral infection (Bestle et al., 2020).

In our study, we present a human-specific *in vitro*, alveolus-on-a-chip model composed of cells of human origin susceptible for a SARS-CoV-2 infection. This model was only recently developed in our lab (Deinhardt-Emmer et al., 2020a). Within the present study, it was modified by using SARS-CoV-2 permissive epithelial cells (Calu-3 cells). The epithelial and vascular (primarily isolated human umbilical vascular endothelial cells; HUVECs) cells were co-cultured with macrophages (primarily isolated peripheral blood mononuclear cells; PBMCs) resembling the human alveolus architecture and function. This composition is not only relevant for the gas exchange but also for an adequate immune response.

We were able to show that SARS-CoV-2 replicates in the epithelial layer while inducing an acute and robust inflammatory response followed by the destruction of the epithelial layer. Interestingly, in this infection scenario, the endothelial cells were not invaded by SARS-CoV-2 and did not propagate the virus, but nevertheless the epithelial/endothelial barrier integrity was disrupted.

## RESULTS

### Efficient SARS-CoV-2 isolation from patients and propagation in cell-culture

To gain fully infectious viral particles for our studies, we collected three respiratory specimens from qRT-PCR-proven COVID-19 patients and performed SARS-CoV-2 propagation in cell culture systems (Vero-76 cells). By repeated infection of host cells and viral replication, we were able to isolate high viral titers originating from three different patients. Within our studies, the SARS-CoV-2 isolates SARS-CoV-2/hu/Germany/Jena-vi005159/2020 (5159), SARS-CoV-2/hu/Germany/Jena-vi005187/2020 (5587) and SARS-CoV-2/hu/Germany/Jena-vi005588/2020 (5588) were employed. Sequencing of virus isolates verified that all three viral strains belong to SARS-CoV-2 (species *Severe acute respiratory syndrome-related coronavirus*, genus *Betacoronavirus*) (Gorbalenya et al., 2020). Phylogenetic analysis revealed a close relationship of SARS-CoV-2 to the SARS-related coronaviruses RaTG13, bat-SL-CoVZXC21 and bat-SL-CoVZC45 (Figure 1A). Within the SARS-CoV-2 clade, the sequences of strains 5587 and 5588 exhibit two base substitutions T8,782C (**nsp1ab:** synonymous) and C28,144T (**nsp8:** S84L), which are characteristic of the all strains of lineage L ((Tang et al., 2020) nomenclature) or lineage B ((Rambaut et al., 2020) nomenclature). Accordingly, 5587 and 5588 clustered with lineage L/lineage B strains in the phylogenetic analysis (Figure 1B). Furthermore, both strains exhibit deletion of **nsp1ab** D448 and two synonymous substitutions (T514C, C5512T). Beside the **nsp8** S84L substitution, strain 5159 has accumulated three additional amino acid substitutions (**S:** D614G, **nsp1ab:** P4715L and **N:** R203K/G204R) which place this virus in lineage B.1.1 according to the proposed SARS-CoV-2 nomenclature of Rambaut et al. (2020) (Figure 1B).

**Figure 1:**
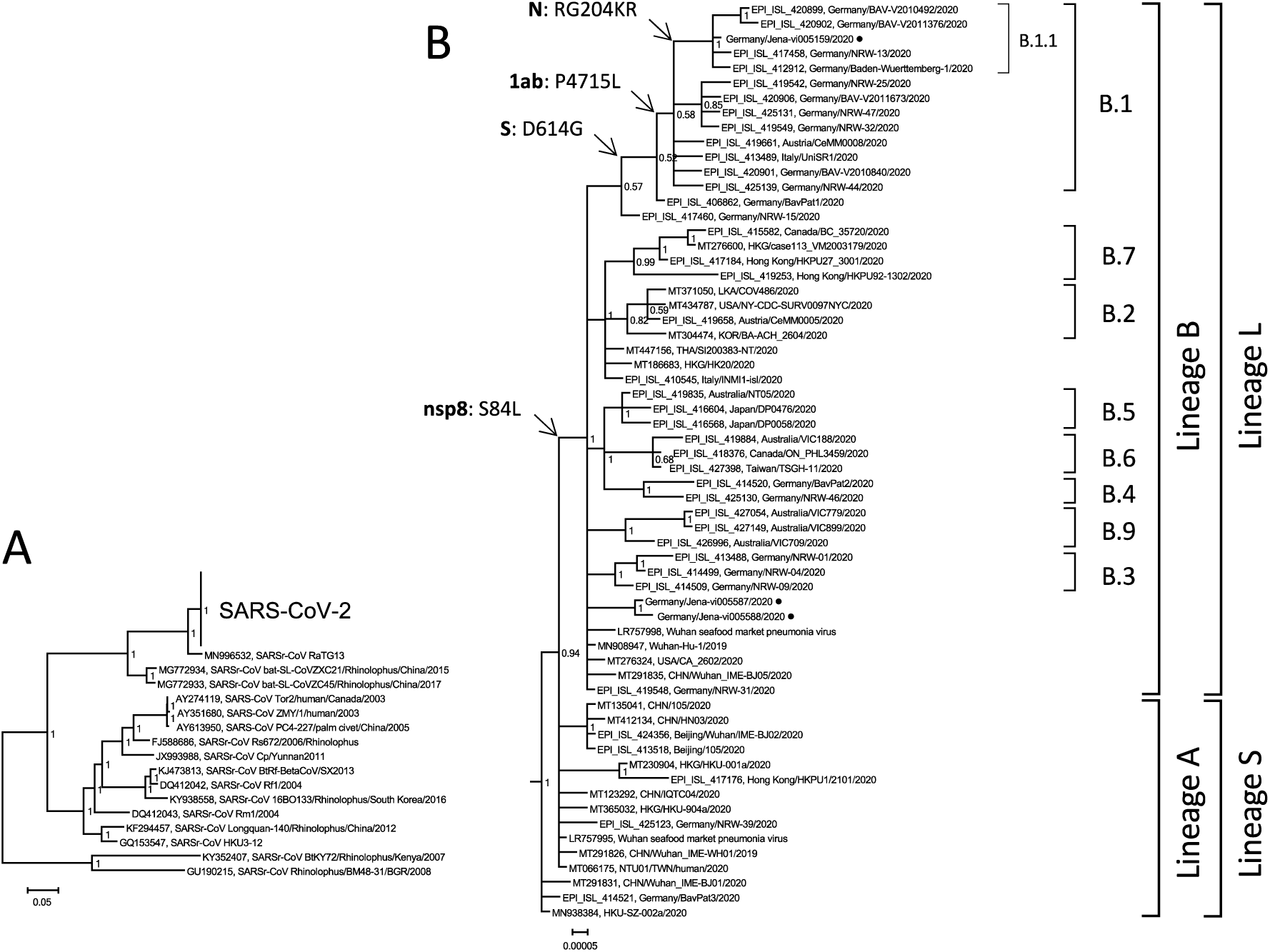
Phylogenetic tree of SARS-CoV-2. (A) Phylogenetic analysis revealed a close relationship of SARS-CoV-2 to the SARS-related coronaviruses RaTG13, bat-SL-CoVZXC21 and bat-SL-CoVZC45. Sequences of strains 5587 and 5588 exhibit two base substitutions T8,782C (**nsp1ab:** synonymous) and C28,144T (**nsp8:** S84L). (B) Accordingly, 5587 and 5588 clustered with lineage L/lineage B strains in the phylogenetic analysis. Both strains exhibit deletion of **nsp1ab** D448 and two synonymous substitutions (T514C, C5512T). Beside the **nsp8** S84L substitution, strain 5159 has accumulated three additional amino acid substitutions (**S:** D614G, **nsp1ab:** P4715L and **N:** R203K/G204R) which place this virus in lineage B.1.1.

### Mono-culture cells can be infected by SARS-CoV-2 and produce replication complexes at ER-derived membranes

At first, we infected mono-cell culture systems with SARS-CoV-2 and compared the infection rate between Vero-76 cells and Calu-3 cells. It is already well known that Vero-76 cells can be efficiently infected by SARS-CoV-2 (Hoffmann et al., 2020; Shang et al., 2020). Using transmission electron microscopy (TEM), we could demonstrate that Vero-76 cells host and efficiently propagate the virus (Figure 2A). Figure 2A (upper panel) illustrates infected Vero-76 cell containing viral replication organelles. In the lower-left panel, protein accumulation and generation of double-membrane vesicles are visible. In the middle panel, virion assembly in the ER–Golgi-intermediate compartment (ERGIC) and a Golgi complex are imaged containing morphologically complete viral particles. Here, the particles are packed to be transported to the cellular surface for virus release, which is demonstrated in the right panel. Some particles are still attached to the host cell membrane, whereas some viral particles are already fully released. These results indicate that SARS-CoV-2 induces replication complexes at ER-derived membranes, which were already shown for other types of coronaviruses (Stertz et al., 2007) and also confirm the findings for Vero E6 cells (Ogando et al., 2020). To better mimic the situation in the human pulmonary alveoli, we performed the infection in Calu-3 cells. In Figure 2B, immunofluorescence measurements compare infected Vero-76 cells with infected Calu-3 cells. In both cell types, viral particles can be visualized to a similar extent using specific antibodies against SARS-CoV-2 spike proteins. These results are confirmed by western blot analysis, demonstrating increased levels of SARS-CoV-2 spike protein in the cell lysates after 8h and 24h (Figure 2C).

**Figure 2:**
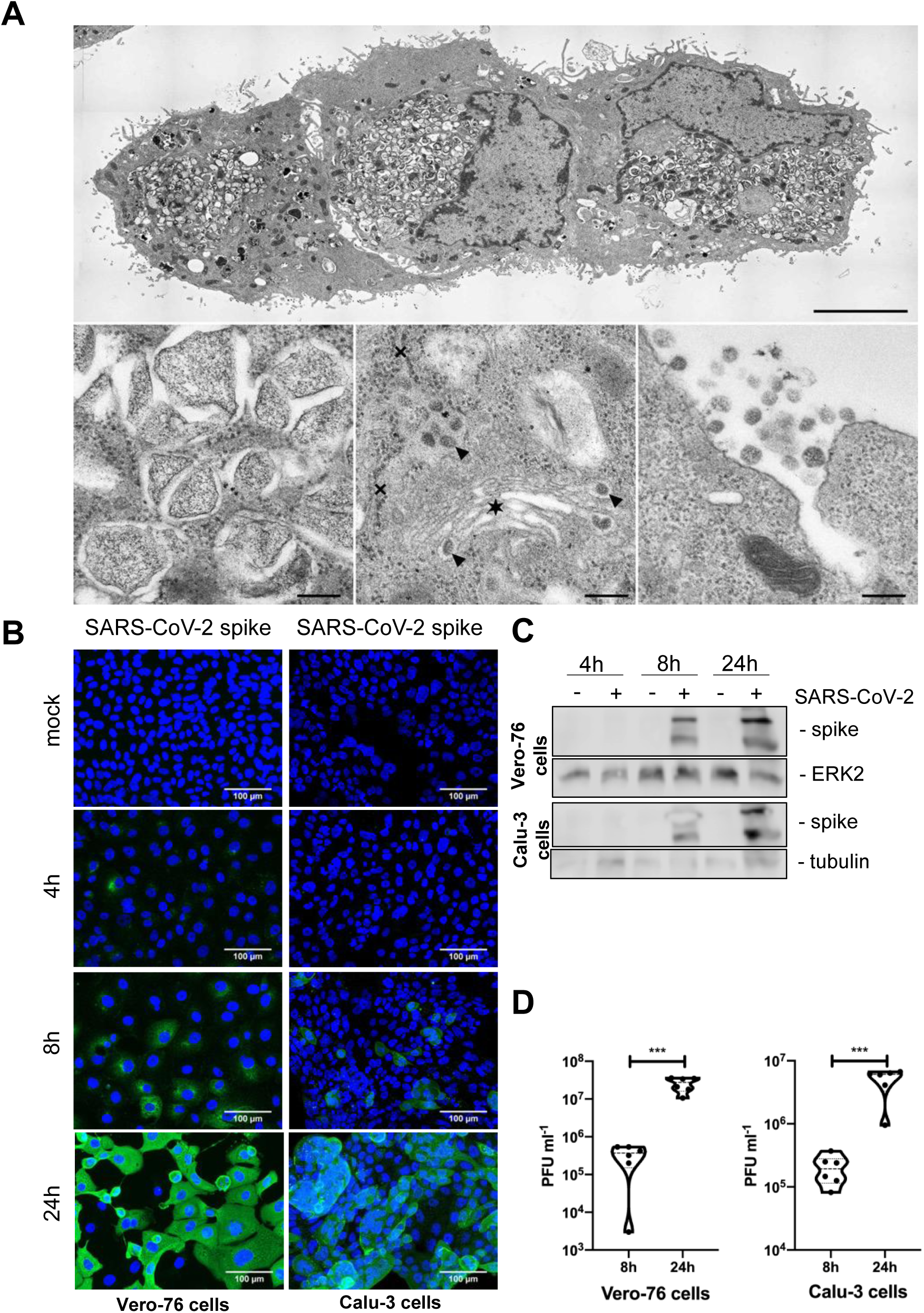
SARS-CoV-2 replicates in Vero-76 and Calu-3 cells. Vero-76 (A-D) and Calu-3 (B-D) cells were left uninfected (mock) (B-D) or were infected (A-D) with a SARS-CoV-2 patient isolate (5159) (MOI=1). (A) Transmission electron microscopy was performed 24h post infection (p.i.): (upper panel, scale bar: 5 μm) overview of 3 SARS-CoV-2-infected Vero-76 cells; (lower left panel, scale bar: 200 nm) generation of double membrane vesicles; (lower middle panel, scale bar: 200 nm) virion assembly in the ER–Golgi-intermediate compartment (ERGIC); (lower right panel, scale bar: 200 nm) viral release. (B) SARS-CoV-2 was visualized by detection of the spike protein via a spike-specific antibody and an Alexa Fluor™ 488-conjugated goat anti-mouse IgG (green). The nuclei were stained with Hoechst 33342 (blue). Immunofluorescence (IF) microscopy was acquired by use of the Axio Observer.Z1 (Zeiss) with a 200×magnification. (C) Total cell lysates were harvested at the times indicated and expression of the spike protein was analyzed by western-blot assay. ERK2 served as loading control. (D) Progeny virus particles were measured in the supernatant by standard plaque assay at the indicated times post infection. Shown are means (±SD) of plaque forming units (PFU) ml^-1^ of three independent experiments including two biological samples. Statistical significance was analyzed by unpaired, two-tailed t-test (***p < 0.001).

Additionally, we could verify progeny virus particles by performing plaque assays from supernatants of both cell types indicating increased replication during ongoing infection (Figure 2D). Measuring viral RNA-loads in different infected host cell types, we found high RNA levels in Vero-76 and Calu-3 cells (Figure S1A). By contrast, HUVEC mono-cell cultures could not be infected by SARS-CoV-2 (Figure S1B).

We further analyzed the host response to the infection by measuring the cytokine mRNA expression of Calu-3 cells. 24h post-infection, many inflammatory cytokines were significantly increased compared to control cells (Figure 3). These results reflect the high cytokine levels found in COVID-19 patients (Costela-Ruiz et al., 2020), indicating that infected epithelial cells contribute to the “cytokine storm” in severe COVID-19 cases. Since different cytokines and chemokines are involved in the infection process, a robust immune response has been associated with a severe clinical course (Coperchini et al., 2020). Our results clearly show that the epithelial cell line Calu-3 can be efficiently infected by SARS-CoV-2, propagates the virus and answer to the viral infection with a strong cytokine release.

**Figure 3:**
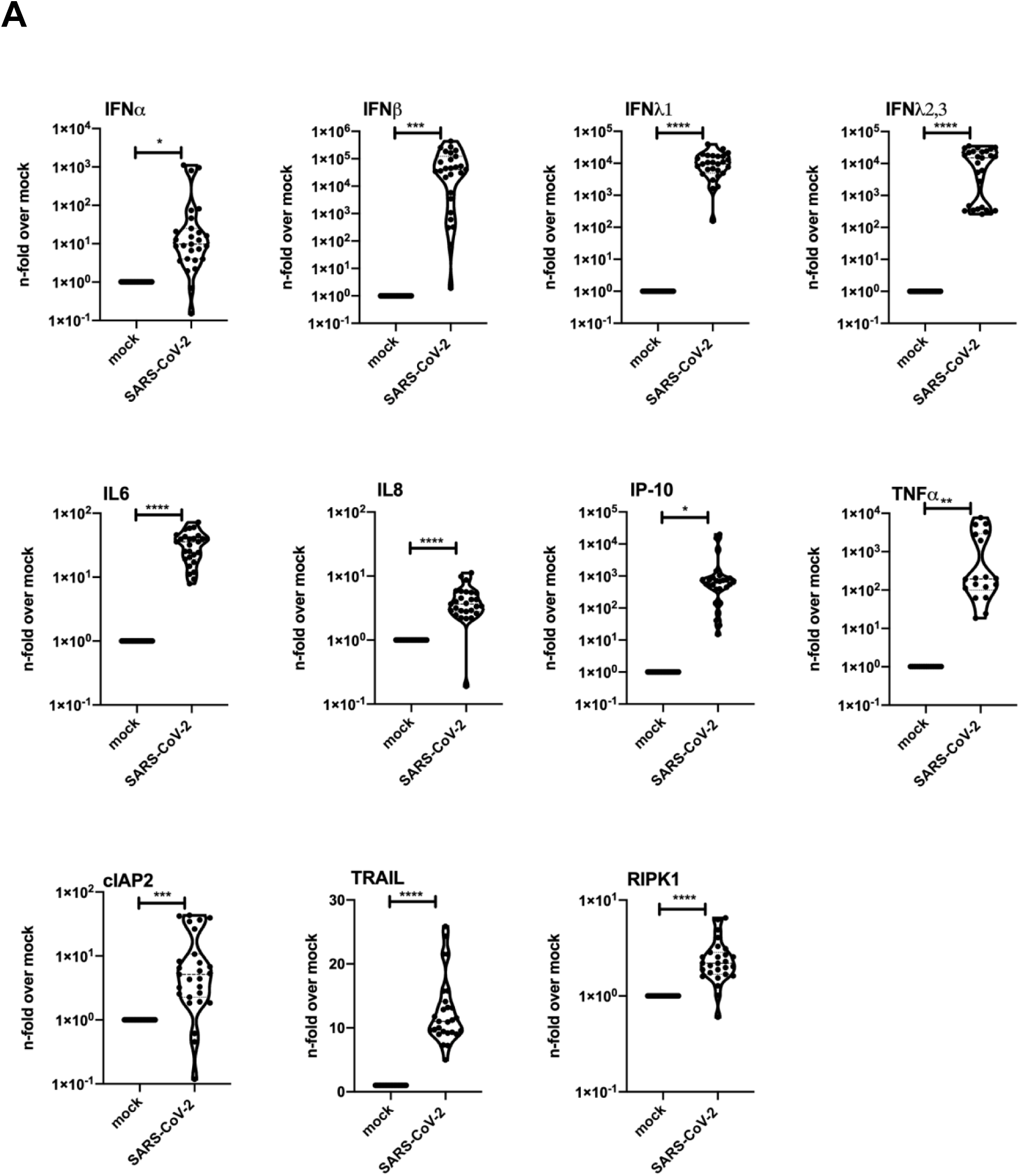
SARS-CoV-2 infection results in induction of antiviral and proinflammatory mRNA synthesis. Calu-3 cells were left uninfected (mock) or were infected with a SARS-CoV-2 patient isolate (5159, 5587, 5588) (MOI=1). RNA-lysates were performed 24h p.i. Levels of IFNα, IFNβ, IFNλ1, IFNλ2,3, IL6, IL8, IP10, TNFα, cIAP2, TRAIL, and RIPK1 Mrna were measured of three patient isolate (5159, 5587, 5588) and two technical samples in 3 independent experiments. Means ± SD of three independent experiments are shown. Levels of mock-treated samples were arbitrarily set as 1. After normalization, two-tailed unpaired t-tests were performed for comparison of mock-treated and SARS-CoV-2-infected and samples. (*p < 0.05, **p < 0.01, ***p < 0.001, ****p < 0.0001).

### SARS-CoV-2 infects epithelial cells within the alveolus-on-a-chip model causing a strong IFN-response

In the next step, we modified our alveolar-on-a-chip model (Deinhardt-Emmer et al., 2020a) by seeding Calu-3 cells on the epithelial side, primarily isolated HUVECs on the endothelial side and integrated PBMCs to represent the immune response. However, macrophages not show an productive viral replication (Yip et al., 2014), they are mainly involved in inflammatory response (Kumar et al., 2020).

The applied system is ventilated and perfused and can be infected with SARS-CoV-2 via the epithelial side. Using the viral particles isolated from the three COVID-19 patients’ specimens, an infection by SARS-CoV-2 on the epithelial cells was proven and the viral particles were propagated (Figure 4A). This effect was still visible after 40h post-infection (Figure S2A). In response to the infection the epithelial cells reacted with a robust cytokine response, demonstrated by elevated IFN-levels in the cell culture supernatants of the epithelial side (Figure 4C).

**Figure 4:**
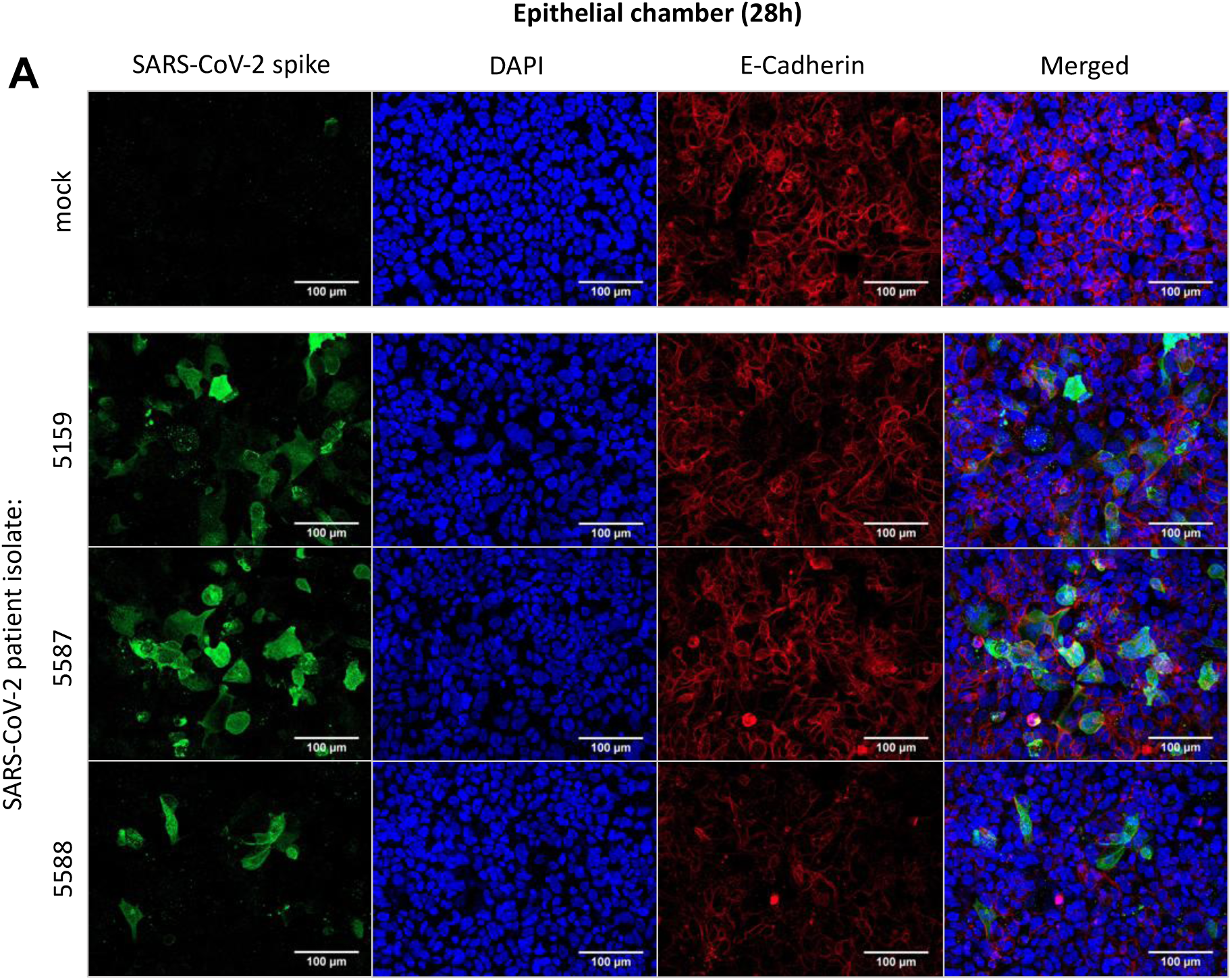
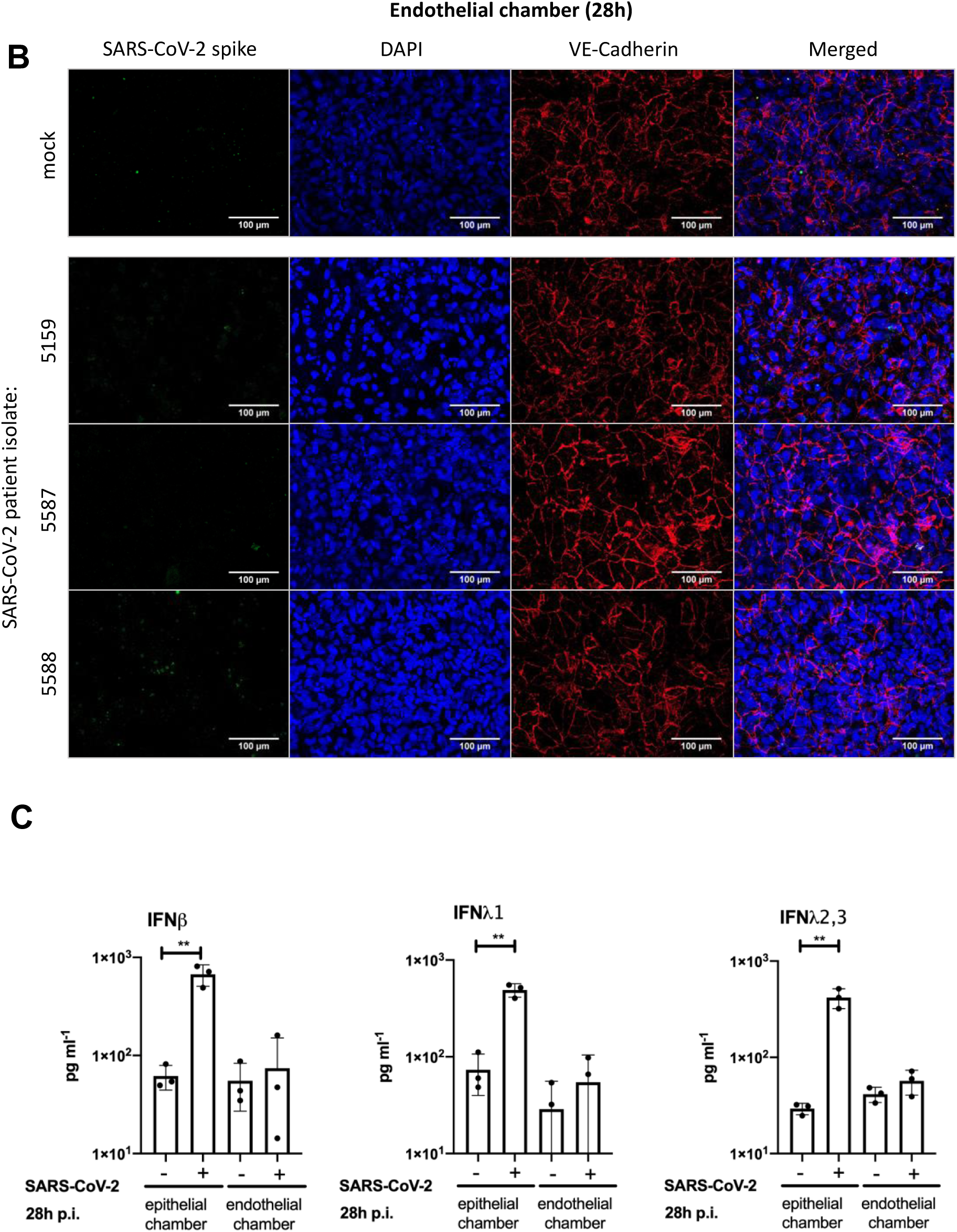
SARS-CoV-2 efficiently infects epithelial cells of the human-alveolus-on-a chip model and provokes type I and III interferon production. (A-C) The epithelial chamber of the alveolus-on-a-chip model was left uninfected (mock) or infected with three different SARS-CoV-2 patient isolates (5159, 5587, 5588) (MOI=1). (A, B) Immunofluorescence staining was performed 28h p.i. and analyzed by immunofluorescence microscopy (Axio Observer.Z1 (Zeiss)). (A) The E-cadherin of the epithelial layer and the (B) VE-cadherin of the endothelial layer were visualized by an anti-E-Cadherin-specific antibody or an anti-VE-Cadherin antiserum, respectively, and a Cy5 goat anti-rabbit IgG (red). (A, B) The SARS-CoV-2 was visualized by detection of the spike protein via a spike-specific antibody and an Alexa Fluor™ 488-conjugated goat anti-mouse IgG (green). The nuclei were stained with Hoechst 33342 (blue). Scale bars represent 100 μm. (C) Production of antiviral cytokines derived from the epithelial side was determined by use of Legendplex Panel (Biolegend, CA, USA). SARS-CoV-2 induced IFNβ, IFNλ1 and IFNλ2,3 release (pg/ml) was measured. Means ± SD of three independent experiments each infected with another patient isolate (5159, 5587, 5588) are shown. Levels of mock-treated samples were arbitrarily set as 1. After normalization, two-tailed unpaired t-tests were performed for comparison of mock-treated and SARS-CoV-2-infected and samples. (**p < 0.01).

In general, the production of IFN is the most efficient way of fighting viral infections; e.g. secretion of type I IFN (IFN-α/β) exhibits direct antiviral effects by inhibiting viral replication (Thiel and Weber, 2008) among many other interferon effects that promote the immune response to infection (Kindler 2016). Yet, evasion strategies for different types of coronaviruses have been described. The viruses express factors and posses strategies to inhibit IFN induction/expression (Thoms et al., 2020) or IFN signaling or to increase IFN resistance, which is reviewed by E. Kindler et al. (2016). Consequently, SARS-CoV-2 is apparently able to cope with the interferon response of epithelial cells, which is reflected by our measurements, demonstrating efficient viral replication and persistence for up to 40h despite a strong epithelial interferon response (Figure 3 and 4).

By contrast, we did not detect viral propagation and did not measure an interferon response at the endothelial side of the biochip (Figure 4C). We could not visualize viral components within endothelial cells neither at 28h (Figure 4B) nor at 40h post-infection (Figure S2B) demonstrating that the viral particles do not productively infect endothelial cells in the human alveolus-on-a-chip model. Additionally, endothelial mono-cell culture systems could not be infected by SARS-CoV-2 (Figure S1A, B) confirming the cell-type specificity of the viral pathogens for lung epithelial cells. This is in line with *in vivo* studies that describe only a weak IFN-response in the serum of COVID-patients (Hu et al., 2020). In animal models with mouse-adapted SARS virus a delayed onset of the IFN-response resulting in immune dysregulation was described (Channappanavar et al., 2016). The weak and delayed IFN-levels in the serum are probably due to the host cell specificity of SARS-CoV-2, as lung epithelium represents the main infection focus and endothelial cells are only hardly/not infected.

Taken together, these results indicate that endothelial cells of the lung model are not the primary target cells of SARS-CoV-2 which is in agreement with previous studies (Bar-On et al., 2020). Further, it is in line with the observation that endothelial layer of the alveolar capillaries of deceased COVID-19 patients were still intact but epithelial tissue was found seriously damaged (Deinhardt-Emmer et al., 2020b). Although the mechanism is not clear the increased proinflammatory cytokine release might cause an endothelial dysfunction. In addition to the origin of endothelial cells from different organs, comorbidities like obesity and diabetes might render endothelial cells susceptible to be infected as recently described (Huertas et al., 2020; Pons et al., 2020). There is clinical evidence for severe courses of COVID-19 in particular, when preexisting endothelial damage can be suspected (Varga et al., 2020).

Further studies are required to elaborate the impact of the endothelial phenotype and the infection conditions at which these cells are targeted by SARS-CoV-2. In the model used in this study, the pneumotropic features of SARS-CoV-2 could be confirmed based on viral uptake and replication in epithelial cells accompanied with an interferon response described in previous studies. The viral particles were not transferred to the neighboring endothelial layer, although the cells were co-cultivated in a bioinspired manner to recreate the alveolar structure.

### SARS-CoV-2 disrupts the alveolar barrier within the alveolus-on-a-chip model

Next, we analyzed the barrier function of the epithelial-endothelial cell layers in our biochip model. Many clinical case reports and studies describe that critical ill COVID-patients develop severe lung destructions and a systemic sepsis-like syndrome (Ackermann et al., 2020; Gao et al., 2020) that could be partly explained by a disrupted barrier function in the lung. From the immunofluorescence image (Figure 4), we could observe some destruction of the epithelial or endothelial layer, in particular 40h post-infection (Figure S2). To better visualize the cell-layers during the infection, we performed scanning electron microscopy (SEM) analysis of the surface structures following 28h post-infection. On the epithelial side we found dead cells and remnants attached to the cell layer. Dying cells are identified by shrinking, balling, disruption of the plasma membrane, and the loss of microvilli, which can be observed in the upper and middle panel of the infected cells, but to a much lesser extent in mock-treated cells (Figure 5A, upper panel). Interestingly, at a higher magnification we could display the viral particles on the surface of the dead cells (Figure 5A, lower panel).

**Figure 5:**
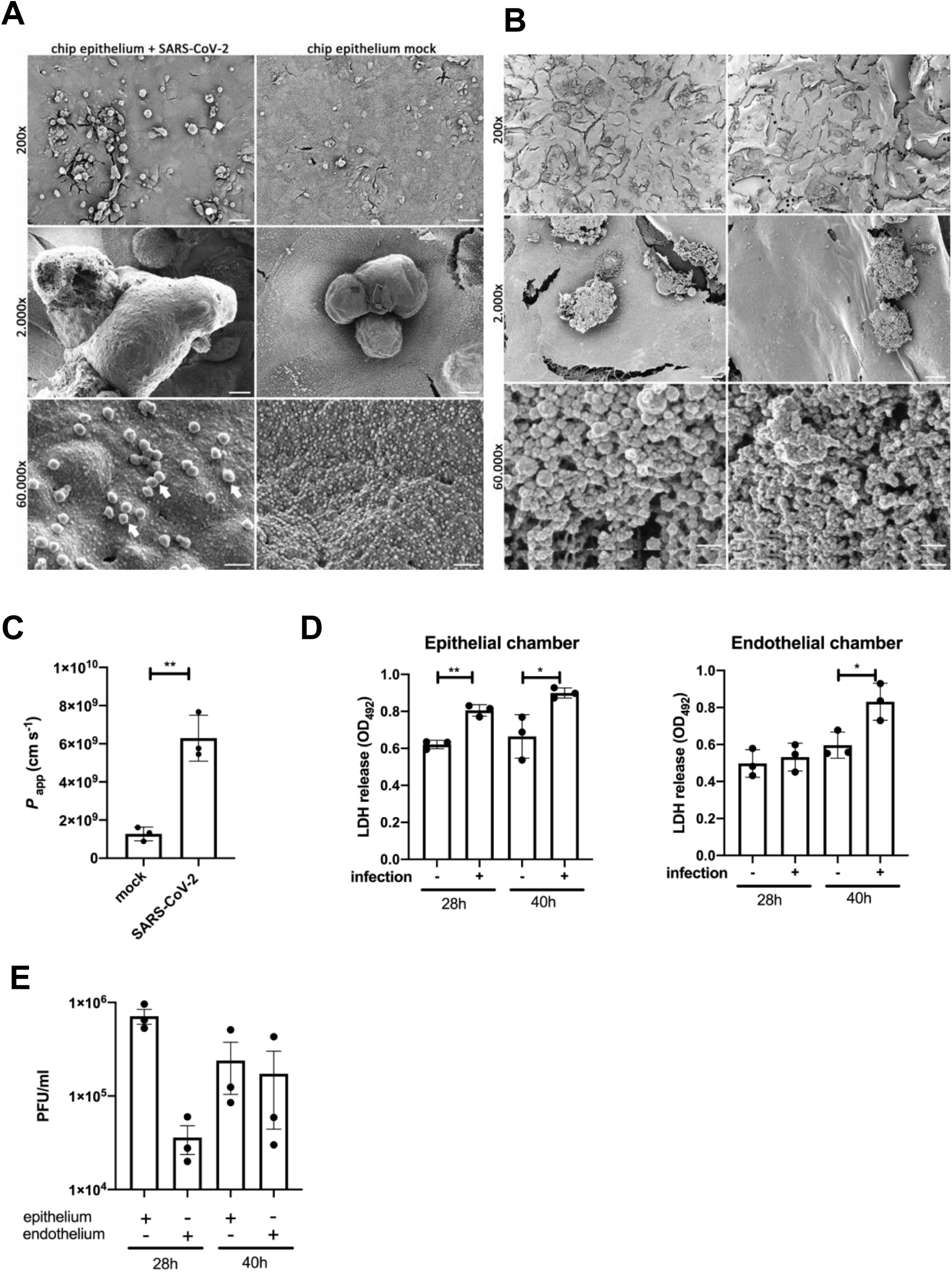
SARS-CoV-2 infection results in the disruption of barrier integrity in the human-alveolus-on-a chip model. The epithelial side of the alveolus-on-a-chip model was left uninfected (mock) or infected with the SARS-CoV-2 patient isolate (5159) (MOI=1) for 28h. An overview (upper panel) of the (A) epithelial layer and (B) endothelial layer are depicted. Dead cells (middle panel) are focused. The surface of dead cells (lower panel) shows particles (arrows) attached to the plasma membranes of the epithelial cells only. Scale bars represent 50 μm (200×magnification), 5 μm (2.000×magnification) and 200 nm (60.000×magnification). (C) Barrier function of the human alveolus-on-a-chip model was analyzed by a permeability assay of mock-infected and SARS-CoV-2-infected human alveolus-on-a-chip model using FITC-dextran at 28h p.i., FITC-dextran was measured via the fluorescence intensity (exc. 488nm; em. 518 nm) and depicted as the permeability coefficient (*P*_*app*_), calculated according to *P*_*app*_ (cm s^-1^) = (dQ/d*t*) (1/AC_o_). Results show significant higher barrier permeability after SARS-CoV-2 infection. (D) Supernatants of the epithelial- and endothelial side of SARS-CoV-2 infected human alveolus-on-a-chip models were used to perform LDH-assays indicating cell membrane rupture at 28h and 40h p.i.. (E) Progeny virus titers were analyzed in the supernatants of the epithelial- and endothelial layer by standard plaque assay. Shown are means (±SD) of (C) three independent experiments each infected with another patient isolate (5159, 5587, 5588), (D) LDH release, and (E) plaque forming units (PFU/ml). Statistical significance was analyzed by unpaired, two-tailed t-test (*p < 0.05, **p < 0.01).

It is well known that respiratory viral pathogens induce cell death (including apoptosis) in the respiratory epithelium, such as the influenza virus (Atkin-Smith et al., 2018). Already the 2003 SARS-CoV led to apoptotic cell death induced by membrane proteins via modulation of the Akt-pathway (Chan et al., 2007). Additionally, prolonged stress of the endoplasmic reticulum (ER) was identified as a trigger for apoptosis (Fung and Liu, 2014). Within the recent pandemic, it has been shown that the largest unique open reading frame (ORF) of the SARS-CoV-2 genome ORF3a is associated with a pro-apoptotic activity (Ren et al., 2020). These studies indicate that the induction of an apoptotic process in the course of SARS-CoV2 infection is highly probable.

By contrast, on the endothelial side, we could not observe differences in the morphological appearance between SARS-CoV-2-infected and mock-treated cells. Here, the cell integrity of the cell layers appeared intact apart from shrinking artefacts due to the drying procedure (Figure 5B). At the high magnification (lower panel) dead cell remnants show granular residues of the cytoplasm indicating the loss of the plasma membrane, but no viral particles could be visualized.

To measure the epithelial and endothelial cell viability we performed LDH-assays on cells grown in the biochip. 28h post-infection we observed in the infected epithelial layer a significantly enhanced release of LDH, whereas SARS-CoV-2 did not induce an enhanced LDH release in endothelial cells (Figure 5D). However, after more extended infection periods (40h; Figure 5D), the barrier function on the endothelial side was also affected. These results suggest that even if the endothelial layer is not infected the cell integrity gets disturbed, most likely by cytokines released by macrophages. In this respect, it is known that high cytokine/interferon levels can induce the disruption of the alveolar barrier function (Broggi et al., 2020; Gustafson et al., 2020; Pelaia et al., 2020).

To further analyze the barrier integrity of the biochip system on a functional level, we performed permeability assays using FITC-dextran to investigate the endothelial and epithelial cell barrier integrity. We were able to show that SARS-CoV-2 significantly increased the tissue permeability with its barrier function severely impaired (Figure 5C). As a consequence of the disrupted barrier, we detected viral particles by performing plaque assays of the cell supernatants in the endothelial chamber at the late time point of 40h (Figure 5E). These results show that endothelial cells are affected by the viral infection at late time points and that the disturbed cell integrity results in translocation of the viral particles over the alveolar barrier.

## DISCUSSION

In this manuscript, we present an *in vitro* human alveolus-on-a-chip model based on human cells that closely mimics alveolar structures and can be efficiently infected by SARS-CoV-2. The epithelial of the alveolar models was demonstrated to be prone to SARS-CoV2 infection and to propagate viral replication with high viral loads. These findings are in line with clinical observations that in lung tissue by far, the highest viral burdens are measured (Carsana et al., 2020). This phenomenon can be explained by the cell tropism of SARS-CoV-2 to airway cells that contribute to the high shedding of viral particles in the respiratory system and the high infectivity of patients via aerosols. Nevertheless, due to the systemic symptoms in severe COVID-patients, it has been discussed, whether other cell types besides the airway epithelium, are targeted by SARS-COV-2, as well. In particular, vascular complications, such as thrombotic events (Helms et al., 2020), could result from the dissemination and propagation of viral particles in the endothelial system. In our model system, we could not confirm viral invasion into endothelial cells, although the cells were cultured in close proximity to the infected epithelium.

Yet, with increased time of infection (40h), the endothelial cells become damaged resulting in a decline in tissue barrier function. This effect is most likely mediated by the cytokine release of the infected neighboring epithelium. Cytokine release is known to disturb various cellular functions, such as protein biosynthesis and barrier integrity. Many studies reveal that most severe cases of SARS-CoV-2 infections are not only due to enhanced viral burden, but to a large extent due to aberrant immune responses (Broggi et al., 2020).

In our manuscript, we present an infection model that could be further used to study several aspects during the SARS-CoV-2 infection: (i) At first, the cellular interaction can be analyzed in detail with increasing complexity. Here, the interaction between endothelial and epithelial cells, and the role of different immune cells that can be integrated into the biochip could be elaborated. (ii) Another crucial aspect is preexisting damage, such as diabetic vascular changes or inflammatory foci that may promote a COVID-19 infection. These factors can be mimicked in the biochip model to investigate their impact on infection development. (iii) A third important issue are novel therapeutic agents. Antiviral and anti-inflammatory therapies can be tested in the biochip model to obtain initial results on their mode of action.

Consequently, our biochip model represents a valuable tool to study many aspects during COVID-19 infections.

## MATERIAL AND METHODS

### Virus isolation, propagation and standard plaque-assays

SARS-CoV-2 was isolated from the respiratory specimen of three different patients and named (SARS-CoV-2/hu/Germany/Jena-vi005159/2020 (5159), SARS-CoV-2/hu/Germany/Jena-vi005187/2020 (5587) and SARS-CoV-2/hu/Germany/Jena-vi005188/2020) (5588) (ethic approvement of the Jena University Hospital, no.: 2018-1263) by using Vero-76 cells. For this, cells were washed 12 h after seeding and infected with 200 μl filtered patient sample (sterilized syringe filter, pore size 0,2 μm) under the addition of Panserin 401 (PanBiotech, Germany). After five days, the cytopathic effect was notable. Then, cells were frozen, centrifuged, and clear supernatants were obtained.

To generate well-defined viral stocks, plaque purification procedures were performed. For this, confluent Vero-76 cell cultures were infected with serial dilutions of virus isolates diluted in EMEM for 60 min at 37°C and 5% CO_2_. Thereafter, inoculum was exchanged with 2 ml MEM/BA (medium with 0.2 % BSA) supplemented with 0.9 % agar (Oxoid, Wesel, Germany), 0.01 % DEAE-Dextran (Pharmacia Biotech, Germany) and 0.2% NaHCO_3_ until plaque formation was observed. Single plaques were marked using inverse microscopy. Contents of these plaques were used to infect confluent Vero-76 cell monolayers in T25 flasks. Cells were incubated at 37°C and 5% CO_2_ until pronounced cytopathic effects were visible. Then, cell cultures were frozen again and clear supernatants were obtained. This plaque purification procedure was repeated again. Finally, virus stocks were generated and titrated using plaque assays. For this, Vero-76 cells were seeded in 6-well plates until a 90 % confluency and infected with serial dilutions of the supernatants in PBS/BA (1 mM MgCl2, 0,9 mM CaCl, 0,2 % BSA, 100 U/ml Pen/Strep) for 90 min at 37°C. After aspiration of the inoculum, cells were incubated with 2 ml MEM/BA (medium with 0.2 % BSA) supplemented with 0.9 % agar (Oxoid, Wesel, Germany), 0.01 % DEAE-Dextran (Pharmacia Biotech, Germany) and 0,2% NaHCO3 at 37°C and 5 % CO_2_ for four days. The visualization was performed by the staining with crystal violet solution (0.2 % crystal violet, 20 % ethanol, 3.5 % formaldehyde in water) and the number of infectious particles (plaque-forming units (PFU) ml^-1^) was determined.

### Sequencing and genome reconstruction

Library preparation was performed according to the “nCoV-2019 sequencing protocol” (dx.doi.org/10.17504/protocols.io.bdp7i5rn) from the ARTICnetwork (https://artic.network/ncov-2019). Briefly, viral RNA was isolated for SARS-CoV-2 virus strains 5159, 5587, and 5588 via the QIAmp viral RNA Kit (Qiagen, Hilden, Germany) according to the manufacturers’ guide. The cDNA preparation was performed using the SuperScript IV (Thermofisher), followed by a multiplex PCR to generate overlapping 400 nt amplicons using version 3 of the primer set (https://github.com/artic-network/artic-ncov2019/tree/master/primer_schemes/nCoV-2019/V3). After PCR cleanup, library preparation was performed using the Ligation Sequencing Kit (LSK-109, Oxford Nanopore Technologies) and the Native Barcoding Expansion (EXP-NBD104, native Barcoding Kit (Oxford Nanopore Technologies)). Sequencing was performed on a MinION device using an R.9.4.1 flow cell (Oxford Nanopore Technologies). Basecalling and genome reconstruction was performed using poreCov v.0.2 with the default settings (https://github.com/replikation/poreCov).

### Cell culture and virus infection

Vero-76 cells were cultured in EMEM with HEPES modification, and 5 mM L-Glutamine. Calu-3 cells were cultured in RPMI-1640 supplemented with 10 % fetal calves’ serum (FCS). M199 was purchased from Lonza (Verviers, Belgium), fetal calf serum (FCS), human serum and endothelial growth supplement were from Sigma (Taufkirchen, Germany).

PBMCs were isolated and cultivated as previously described (Deinhardt-Emmer et al., 2020a). Human umbilical vein endothelial cells (HUVEC) were isolated from anonymously acquired human umbilical cords according to the Declaration of Helsinki, “Ethical principles for Medical Research Involving Human Subjects” (1964). After rinsing the cord veins with 0.9 % NaCl, endothelial cells were detached with collagenase (0.01 %, 3 min at 37 °C), suspended in M199/10 % FCS, washed once (500 x g, 6 min) and seeded on a cell culture flask coated with 0.2 % gelatin. 24 h later, full growth medium was added (M199, 17.5 % FCS, 2.5 % human serum, 7.5 μg/ml endothelial mitogen, 7.5 U/ml heparin, 680 μM glutamine, 100 μM vitamin C, 100 U/ml penicillin, 100 μg/ml streptomycin). HUVEC from the second passage were seeded on 30-mm dishes or on 90-mm dishes at a density of 27,500 cells/cm^2^. Experiments were performed 72 h after seeding. For the cultivation of the human alveolus-on-a-chip model we used Calu-3 cells and macrophages at the epithelial side, and HUVECs at the endothelial side. The Multiorgan tissue flow (MOTiF) biochips were manufactured and obtained from microfluid ChipShop GmbH (Jena, Germany), as explained previously (Deinhardt-Emmer et al., 2020a).

For infection of Vero-76 or Calu-3 cells, cells were washed with PBS and either left uninfected (mock) or infected with SARS-CoV-2 with a multiplicity of infection (MOI) of 1 for 120 min in medium (EMEM with HEPES modification and 5 mM L-Glutamine for Vero-76 cells and RPMI 1640 for Calu-3 cells) supplemented with 10 % FCS. Subsequently, supernatants were removed, and cells were supplemented with fresh medium supplemented with 10 % FCS and further incubated for the times indicated at 37°C, 5 % CO_2_.

For the infection of the human alveolus-on-a-chip, cells were washed with PBS once, followed by treatment with PBS (mock) at 37°C and RPMI (0.2 % autologous human serum, 1 mM MgCl_2_, 0.9 mM CaCl_2_) or infection with SARS-CoV-2 virus (1 MOI). After 90 min incubation cells were washed and supplemented with medium. Afterwards, cells were incubated for the indicated times at 37°C, 5 % CO_2_.

### Transmission electron microscopy

Confluent monolayers of Vero-76 cells (9 cm petri dishes) were infected with SARS-CoV-2 (isolate 5159) using an MOI of 1. After 24 h, supernatants were removed, and samples were fixed with freshly prepared modified Karnovsky fixative consisting of 4 % w/v paraformaldehyde and 2.5 % v/v glutaraldehyde in 0.1 M sodium cacodylate buffer pH 7.4 for 1h at room temperature. After washing 3 times for 15 min each with 0.1 M sodium cacodylate buffer (pH 7.4) the cells were post-fixed with 2 % w/v osmium tetroxide for 1h at room temperature. Subsequently, the cells were rewashed with 0.1 M sodium cacodylate buffer (pH 7.4), thoroughly scraped off the petri dishes, and pelleted by centrifuging at 600 x g for 10 min. During the following dehydration in ascending ethanol series, post-staining with 1 % w/v uranyl acetate was performed. Afterwards, the pellets were embedded in epoxy resin (Araldite) and ultrathin sectioned (70 nm) using a Leica Ultracut S (Leica, Wetzlar, Germany). Finally, the sections were mounted on filmed Cu grids, post-stained with lead citrate, and studied in a transmission electron microscope (EM 900, Zeiss, Oberkochen, Germany) at 80 kV and magnifications of 3,000x to 85,000x. For image recording, a 2K slow-scan CCD camera (TRS, Moorenweis, Germany) was used.

### Immunofluorescence microscopy

Membranes of the human alveolus-on-a-chip were fixed for at least 30 min with 4 % paraformaldehyde at 37°C and permeabilized with 0.1 % saponin buffer for one hour at room temperature. For the alveolus-on-a-chip model the membrane was removed from the chip after fixation and before permeabilization and cut in two halves to analyze either the epithelial or the endothelial side. Infection by SARS-CoV-2 was visualized using mouse anti-SARS-CoV-2 spike (GeneTex; #GTX632604) IgG monoclonal, primary antibodies and AlexaFluor® goat anti-mouse IgG polyclonal antibodies (Dianova; # 115-545-146). The nuclei were stained with bisBenzimide H 33342 trihydrochloride (Hoechst 33342) (Merck; #14533). Rabbit anti-E-cadherin IgG monoclonal (CellSignaling; 3195S) or rabbit anti-VE-cadherin polyclonal, primary antibodies (CellSignaling; 2158S) and Cy5 goat anti-mouse IgG polyclonal antibodies (Dianova; #111-175-144) were used to detect cell borders of Calu-3 or HUVEC cells on the membrane of the alveolus-on-a-chip model, respectively. Primary antibodies were added 1:100, overnight at 4°C. Afterwards, secondary antibodies and Hoechst 33342 were added 1:100 and 1:1000 for 1 h, at room temperature and in the dark. Cells and membranes were mounted with fluorescence mounting media (Dako; #S3023).

Images were acquired using an Axio Observer.Z1 microscope (Zeiss) with Plan Apochromat 20x/0.8 objective (Zeiss), ApoTome.2 (Zeiss) and Axiocam 503 mono (Zeiss) and the software Zen 2.6 (blue edition; Zeiss). Apotome defolding with phase error correction and deconvolution was done by the software Zen 2.6 as well. Fiji V 1.52b (ImageJ) was used for further image processing, including Z-stack merging with maximum intensity projection and gamma correction. Parameters were kept the same for all pictures which were compared with each other.

### Scanning electron microscopy

The fixation of the cells was performed inside the human alveolus-on-a-chip model by using the same fixative as for TEM for 60 min at room temperature as described previously (Deinhardt-Emmer et al., 2020a; Maurer et al., 2019; Rennert et al., 2015). Afterwards, the chips were rinsed three times with fresh cacodylate buffer for 10 min each and the membranes were cut out. After post-fixation with 2 % w/v osmiumtetroxide for 1h the samples were dehydrated in ascending ethanol concentrations (30, 50, 70, 90 and 100 %) for 15 min each. Subsequently, the samples were critical-point dried using liquid CO_2_ and sputter coated with gold (thickness approx. 2 nm) using a CCU-010 sputter coater (safematic GmbH, Zizers, Switzerland). The specimens were investigated with a field emission SEM LEO-1530 Gemini (Carl Zeiss NTS GmbH, Oberkochen, Germany).

### Western-Blot Analysis

For western blotting, cells were lysed with Triton lysis buffer (TLB; 20 mM Tris-HCl, pH 7.4; 137 mM NaCl; 10% Glycerol; 1% Triton X-100; 2 mM EDTA; 50 mM sodium glycerophosphate, 20 mM sodium pyrophosphate; 5 μg ml^-1^ aprotinin; 5 μg ml^-1^ leupeptin; 1 mM sodium vanadate and 5 mM benzamidine) for 30 min. Cell lysates were cleared by centrifugation, supplemented with 5x Lämmli buffer (10% SDS, 50% glycerol, 25% 2-mercaptoethanol, 0.02% bromophenol blue, 312 mM Tris 6.8 pH) (diluted 1:5), boiled for 10 min (95°C), and subjected to SDS-PAGE and subsequent blotting. For the detection of SARS-CoV-2 spike protein a rabbit polyclonal anti-SARS-CoV-2 spike S2 antibody (Sino Biological #40590-T62) was used.

### Lactate Dehydrogenase Cytotoxicity Assay

Cell cytotoxicity was determined with CyQUANT Lactate Dehydrogenase (LDH) Cytotoxicity Assay Kit (Invitrogen/Thermo Fisher Scientific, Waltham, USA) according to the manufacturer’s instructions. Cells were infected as previously described. After infection 25 μl of the supernatant was transferred in technical duplicates to a 96-well plate and mixed with 25 μl of the LDH cytotoxicity assay reagent. The plate was incubated at 37°C for 30 min. Stop-Solution (25 μl) was added, and OD_492nm_ was directly measured using a TECAN Spectra fluor plate reader (Tecan Group Ltd, Maennedorf, Switzerland). The OD_620nm_ was subtracted to correct for background signal.

### Permeability Assay

To test the permeability of the epithelial and endothelial barrier, 1 mg ml^-1^ of 3–5 kDa fluorescein isothio-cyanate (FITC)-dextran (Sigma-Aldrich, Germany) in phenol-red free DMEM/F12 medium (Sigma-Aldrich, Germany) was injected into the upper chamber of the chip. The lower chamber contained only phenol red free DMEM/F12. The alveolus model was incubated for 60 min under static conditions. Afterwards, the media from the lower and upper chambers were collected, and fluorescence intensity (exc. 488nm; em. 518 nm) was measured in a 96-well *μ*Clear black plate (Greiner BioOne, Frickenhausen, Germany) by a BMG Labtech FLUOStar Omega microplate reader (BMG Labtech GmbH, Ortenberg, Germany). The permeability coefficient (*P*_*app*_) was calculated according to *P*_*app*_ (cm s^-1^) = (dQ/d*t*) (1/AC_o_). For this, dQ/d*t* represent the steady-state flux (g s^-1^), *A* the culture surface area (cm^2^) and C_o_ the initial concentration (mg ml^-1^) (Thomas et al., 2017).

### Detection of mRNA-expression by using qRT-PCR

For RNA isolation cells were lysed with 350 μl RLT lysis buffer and detached from the plate using a rubber cell scraper. RNA isolation was performed using the RNeasy Mini Kit (QIAGEN, Hilden, Germany) according to the manufacturer’s protocol. RNA concentration was measured using the Nano Drop Spectrophotometer ND-1000 (preqlab/vwr, Radnor, USA).

For cDNA synthesis, the QuantiNova Reverse Transcription Kit (QIAGEN, Venlo, Netherlands) was used. RNA was thawed on ice. 400 nanogram (ng) RNA were diluted in RNase free water to a volume of 13 μl. 2 μl gDNA removal mix were added to the diluted RNA; followed by incubation at 45°C for 2 min. After incubating the samples for at least 1 min on ice, 5 μl of RT master mix (containing 4 μl Reverse Transcription Mix and 1 μl Reverse Transcription enzyme per sample) were added. The resulting mixture was incubated for 3 min at 25°C, followed by incubation at 45°C for 10 min and an inactivation step at 85°C for 5 min. The cDNA was either directly used for the subsequent experiments or stored at −20°C.

qRT-PCRs were performed using the QuantiNova SYBR Green PCR Kit (QIAGEN, Venlo, Netherlands). 1 μl cDNA was added to 19 μl master mix (containing 10 μl SYBR Green, 1.5 μl Forward Primer (10 μM), 1.5 μl Reverse Primer (10 μM) and 6 μL RNase free ddH_2_O per sample; for primer sequences see table 1), and the real time PCR reaction was started using the following cycle conditions: 95l°C for 2 min, followed by 40 cycles of 95°C for 5 sec and 60°C for 10 sec. The qPCR cycle was ended by a stepwise temperature-increase from 60°C to 95°C (1°C every 5 sec).

### Detection of SARS-CoV-2 by using qRT-PCR

For the determination of SARS-CoV-2, we used the QIAamp Viral RNA Mini Kit (Qiagen, Hilden, Germany) according to manufacturer’s guide. A qRT-PCR from RIDAgene (r-biopharm, Darmstadt, Germany) followed on Rotor-Gene Q (Qiagen, Hilden, Germany) to detect the E-gene of SARS-CoV-2. The RNA standard curve, prepared from the positive control of the RIDAgene (r-biopharm, Darmstadt, Germany) kit, Cycle conditions were set as follows: 10 min at 58°C, 1 min at 95°C and 45 cycles of 95°C for 15 sec and 60°C for 30 sec.

### Statistical analysis

Statistical analyses were performed using Prism 8 (GraphPad Software). Statistical methods are described in the figure legends..

## Author contributions

SDE, BL and CE conceived and designed the experiments, SDE, SB, CH, LG, AH, RZ, FH, CB, MM, MS, and SN performed the experiments, SDE, SB, CH, LG, RZ, ASM, CB, SN, BL and CE analyzed the data, SDE, BL and CE wrote the manuscript, SDE, ASM, MWP, RH, SN, BL and CE provided resources. All authors critically read and commented on the manuscript.

## Acknowledgments

We thank the team at the Placenta Laboratory of the Jena University Hospital for supplying umbilical cords for HUVEC isolation and the excellent technical work of Elke Teuscher regarding the preparation. Furthermore, we thank Stefanie Kynast for the excellent cell cultivation and laboratory assistance. The authors also thank Anika Hopf (Center for Electron Microscopy) for her excellent work in sample preparation.

## Figure legends

**Figure S1:**
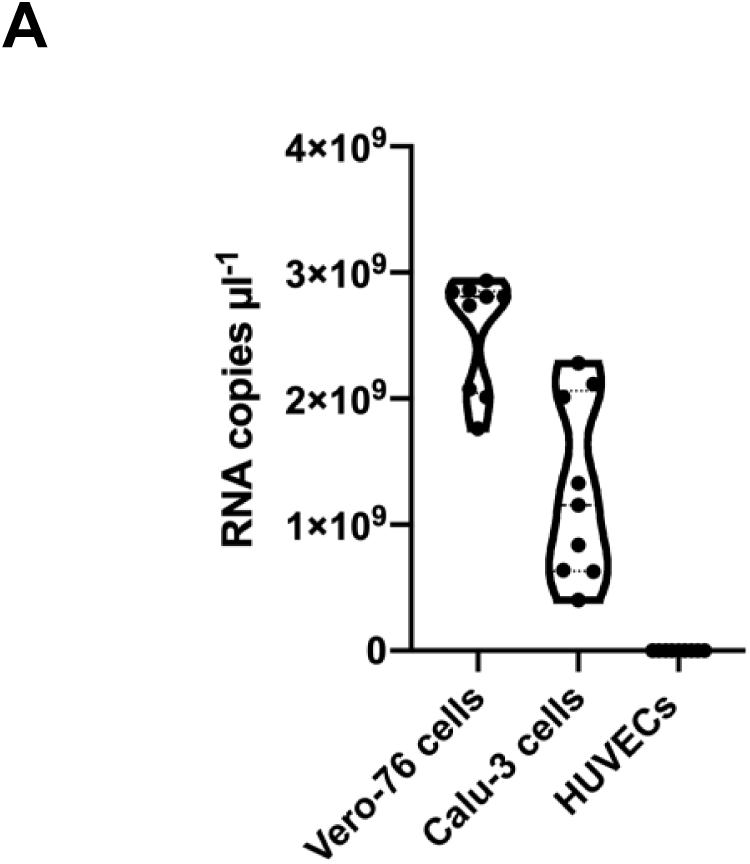
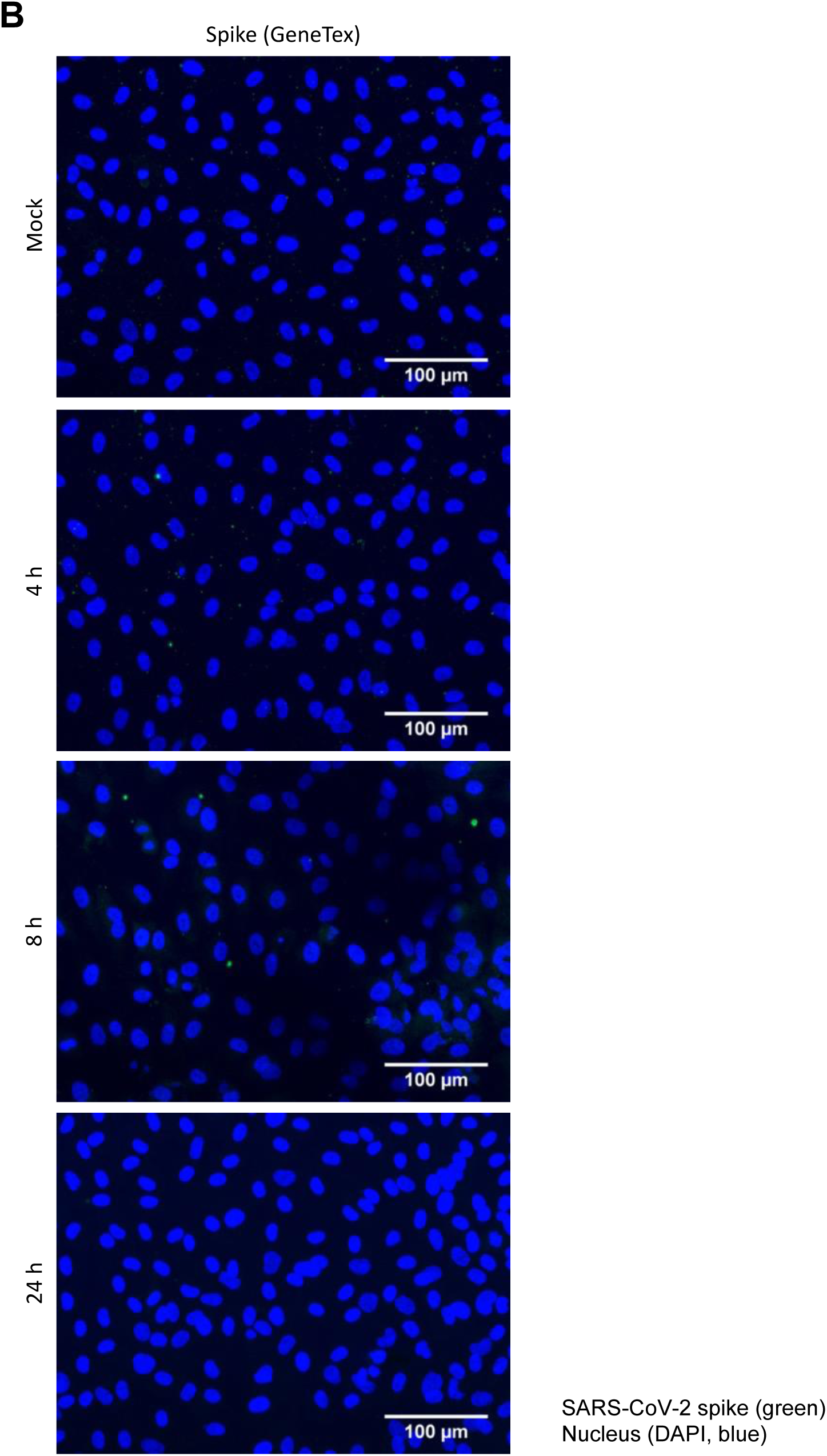
SARS-CoV-2 infects epithelial cells productively. (A) Vero-76, Calu-3, and HUVECs were infected with a SARS-CoV-2 patient isolate (5159, 5587, 5588) (MOI=1). RNA-lysates were performed 24h p.i. and copies of viral RNA (E-gene) were determined by r-biopharm qRT-PCR. Means ± SD of three independent experiments are shown. (B) HUVECs were infected with a SARS-CoV-2 patient isolate (5159) (MOI=1) for 4h, 8h, and 24h. SARS-CoV-2 was visualized by detection of the spike protein via a spike-specific antibody and an Alexa Fluor™ 488-conjugated goat anti-mouse IgG (green). The nuclei were stained with Hoechst 33342 (blue). Immunofluorescence (IF) microscopy was acquired by use of the Axio Observer.Z1 (Zeiss) with a 200×magnification.

**Figure S2:**
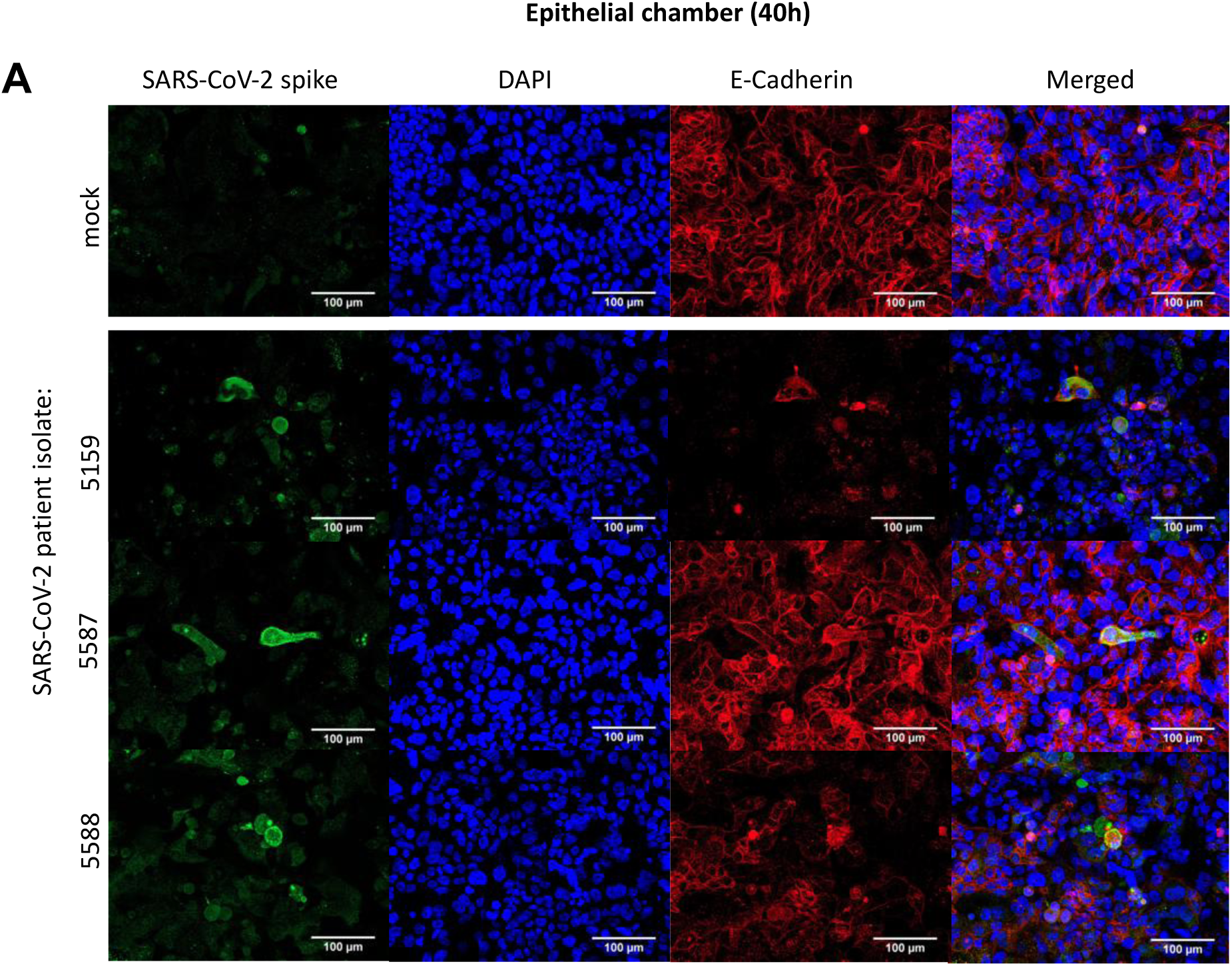
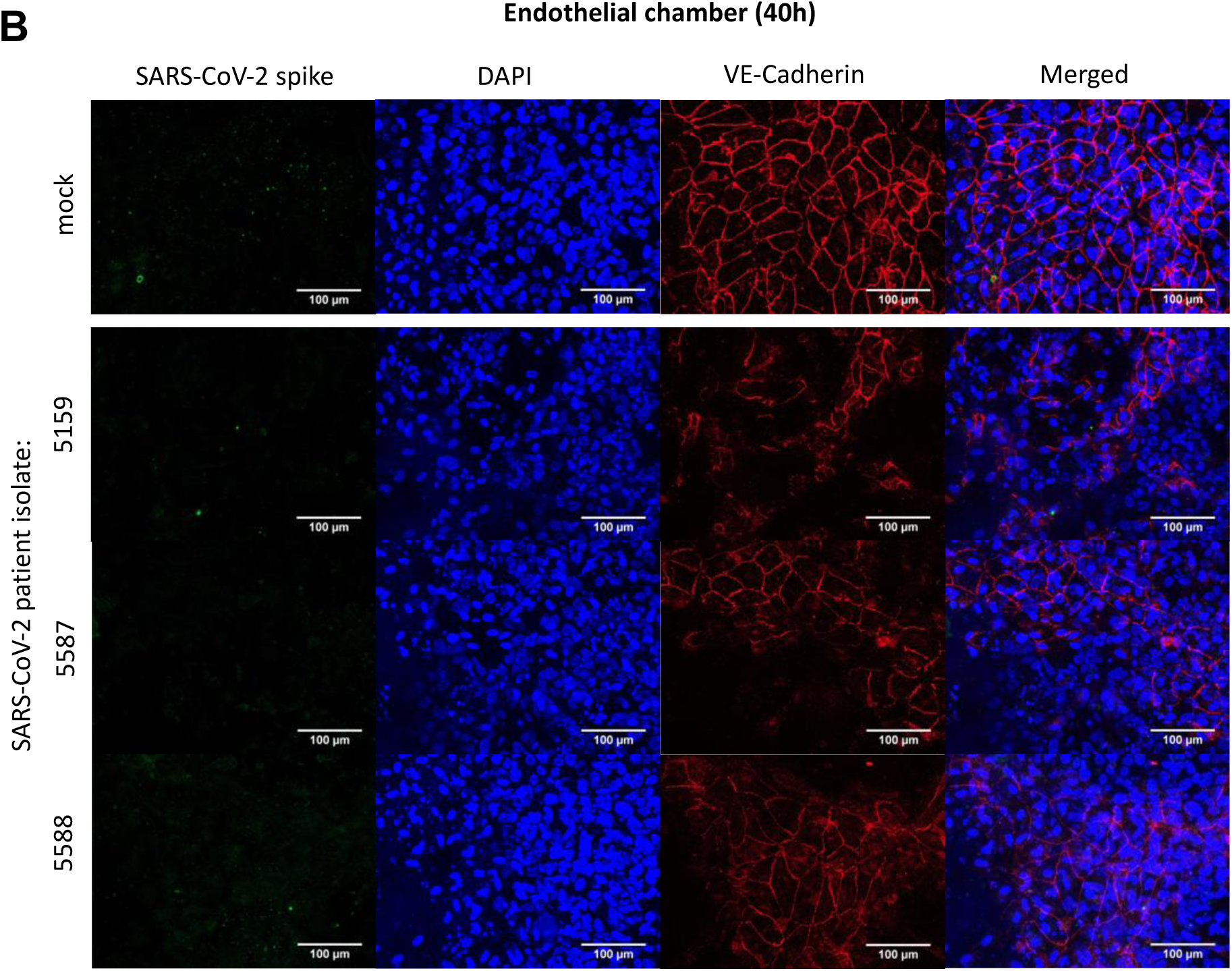
Infection with SARS-CoV-2 results in the disruption of the epithelial- and endothelial barrier. The epithelial side of the alveolus-on-a-chip model was left uninfected (mock) or infected with three different SARS-CoV-2 patient isolates (5159, 5587, 5588) (MOI=1). Immunofluorescence staining was performed 40h p.i., (A) The E-cadherin of the epithelial layer and the (B) VE-cadherin of the endothelial layer were visualized by an anti-E-Cadherin-specific antibody or an anti-VE-Cadherin antiserum, respectively, and a Cy5 goat anti-rabbit IgG (red). (A, B) The SARS-CoV-2 was visualized by detection of the spike protein via a spike-specific antibody and an Alexa Fluor™ 488-conjugated goat anti-mouse IgG (green). The nuclei were stained with Hoechst 33342 (blue). Scale bars represent 100 μm.

## STAR ⋆ METHODS

### KEY RESOURCES TABLES

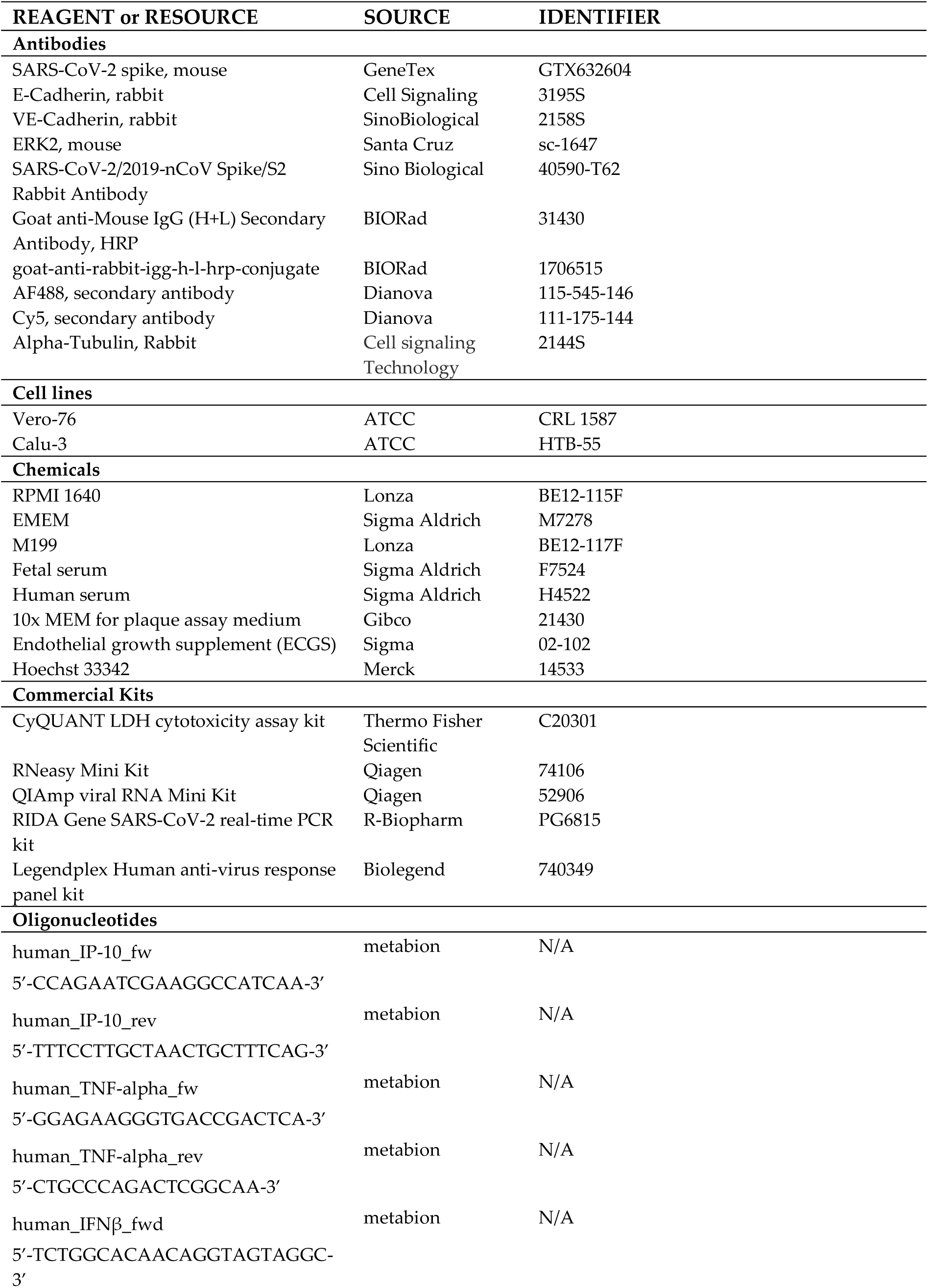
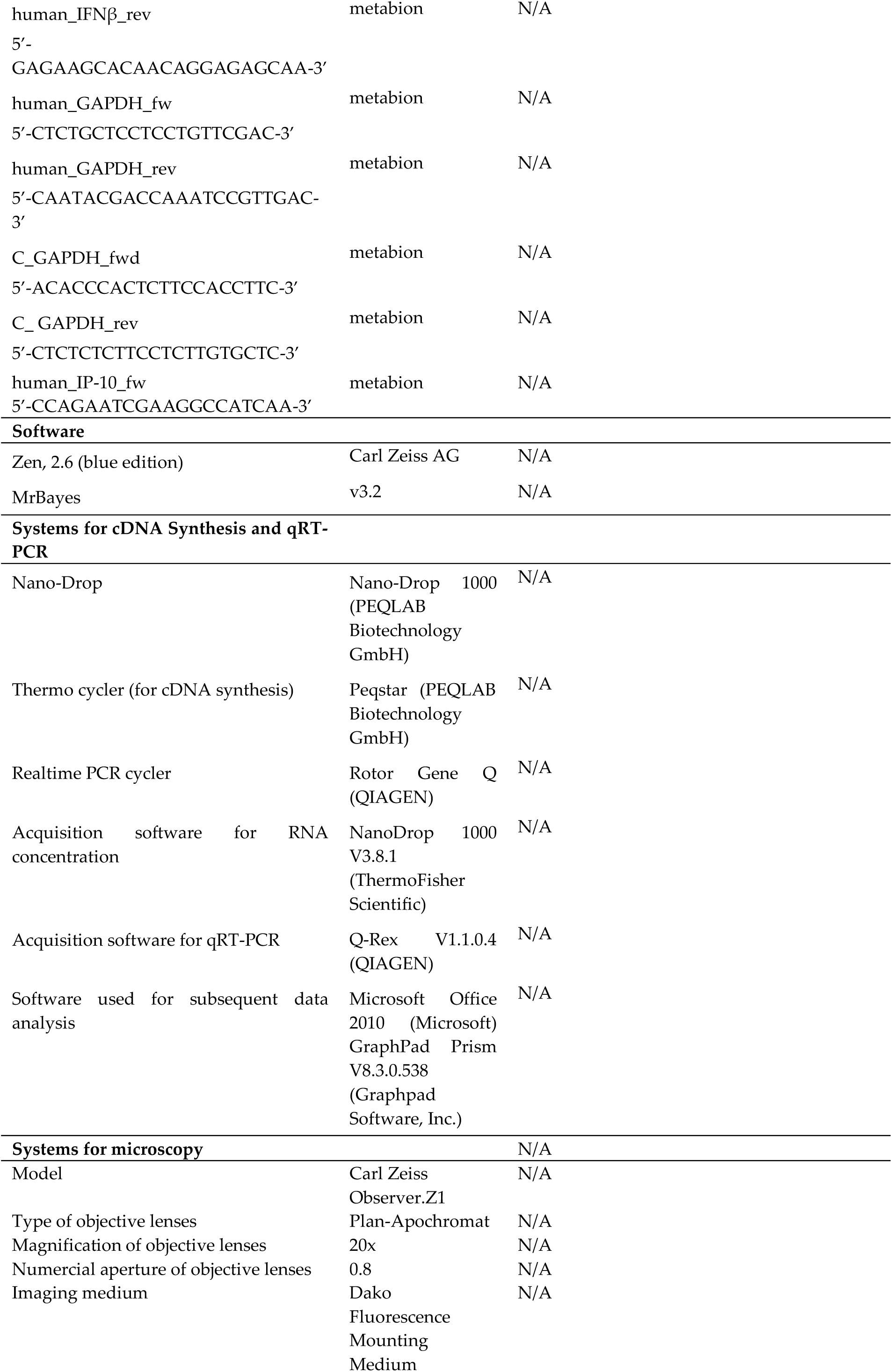
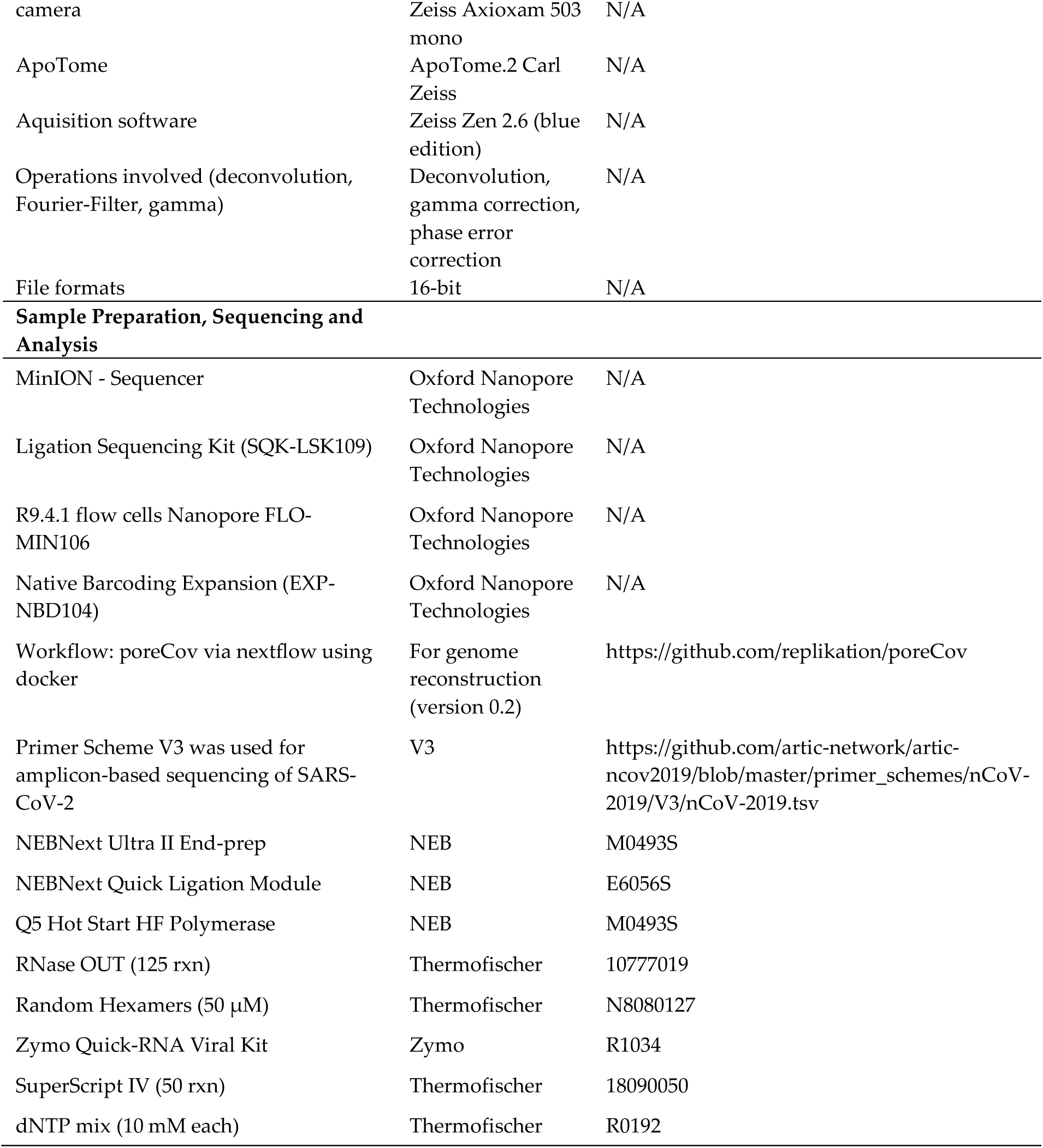

## References

Ackermann, M., S.E. Verleden, M. Kuehnel, A. Haverich, T. Welte, F. Laenger, A. Vanstapel, C. Werlein, H. Stark, A. Tzankov, W.W. Li, V.W. Li, S.J. Mentzer, and D. Jonigk. 2020. Pulmonary Vascular Endothelialitis, Thrombosis, and Angiogenesis in Covid-19. New England Journal of Medicine 383:120–128.

Atkin-Smith, G.K., M. Duan, W. Chen, and I.K.H. Poon. 2018. The induction and consequences of Influenza A virus-induced cell death. Cell Death & Disease 9:1002.

Bar-On, Y.M., A. Flamholz, R. Phillips, and R. Milo. 2020. SARS-CoV-2 (COVID-19) by the numbers. eLife 9:e57309.

Becker, R.C. 2020. COVID-19 update: Covid-19-associated coagulopathy. J Thromb Thrombolysis 50:54–67.

Bestle, D., M.R. Heindl, H. Limburg, T. Van Lam van, O. Pilgram, H. Moulton, D.A. Stein, K. Hardes, M. Eickmann, O. Dolnik, C. Rohde, H.D. Klenk, W. Garten, T. Steinmetzer, and E. Böttcher-Friebertshäuser. 2020. TMPRSS2 and furin are both essential for proteolytic activation of SARS-CoV-2 in human airway cells. Life Sci Alliance 3:

Broggi, A., S. Ghosh, B. Sposito, R. Spreafico, F. Balzarini, A. Lo Cascio, N. Clementi, M. De Santis, N. Mancini, F. Granucci, and I. Zanoni. 2020. Type III interferons disrupt the lung epithelial barrier upon viral recognition. Science (New York, N.Y.)

Carsana, L., A. Sonzogni, A. Nasr, R.S. Rossi, A. Pellegrinelli, P. Zerbi, R. Rech, R. Colombo, S. Antinori, M. Corbellino, M. Galli, E. Catena, A. Tosoni, A. Gianatti, and M. Nebuloni. 2020. Pulmonary post-mortem findings in a series of COVID-19 cases from northern Italy: a two-centre descriptive study. The Lancet Infectious Diseases

Chan, C.-M., C.-W. Ma, W.-Y. Chan, and H.Y.E. Chan. 2007. The SARS-Coronavirus Membrane protein induces apoptosis through modulating the Akt survival pathway. Arch Biochem Biophys 459:197–207.

Channappanavar, R., A.R. Fehr, R. Vijay, M. Mack, J. Zhao, D.K. Meyerholz, and S. Perlman. 2016. Dysregulated Type I Interferon and Inflammatory Monocyte-Macrophage Responses Cause Lethal Pneumonia in SARS-CoV-Infected Mice. Cell Host Microbe 19:181–193.

Coperchini, F., L. Chiovato, L. Croce, F. Magri, and M. Rotondi. 2020. The cytokine storm in COVID-19: An overview of the involvement of the chemokine/chemokine-receptor system. Cytokine Growth Factor Rev 53:25–32.

Costela-Ruiz, V.J., R. Illescas-Montes, J.M. Puerta-Puerta, C. Ruiz, and L. Melguizo-Rodríguez. 2020. SARS-CoV-2 infection: The role of cytokines in COVID-19 disease. Cytokine Growth Factor Rev S1359-6101(1320)30109-X.

Deinhardt-Emmer, S., K. Rennert, E. Schicke, Z. Cseresnyés, M. Windolph, S. Nietzsche, R. Heller, F. Siwczak, K.F. Haupt, S. Carlstedt, M. Schacke, M.T. Figge, C. Ehrhardt, B. Löffler, and A.S. Mosig. 2020a. Co-infection with Staphylococcus aureus after primary influenza virus infection leads to damage of the endothelium in a human alveolus-on-a-chip model. Biofabrication 12:025012.

Deinhardt-Emmer, S., D. Wittschieber, J. Sanft, S. Kleemann, S. Elschner, K.F. Haupt, V. Vau,C. Häring, J. Rödel, A. Henke, C. Ehrhardt, M. Bauer, M. Philipp, N. Gaßler, S. Nietzsche, B. Löffler, and G. Mall. 2020b. Early postmortem mapping of SARS-CoV-2 RNA in patients with COVID-19 and correlation to tissue damage. bioRxiv 2020.2007.2001.182550.

Fung, T.S., and D.X. Liu. 2014. Coronavirus infection, ER stress, apoptosis and innate immunity. Front Microbiol 5:

Gao, Y.M., G. Xu, B. Wang, and B.C. Liu. 2020. Cytokine storm syndrome in coronavirus disease 2019: A narrative review. J Intern Med

George, P.M., A.U. Wells, and R.G. Jenkins. 2020. Pulmonary fibrosis and COVID-19: the potential role for antifibrotic therapy. The Lancet Respiratory Medicine

Gorbalenya, A.E., S.C. Baker, R.S. Baric, R.J. de Groot, C. Drosten, A.A. Gulyaeva, B.L. Haagmans, C. Lauber, A.M. Leontovich, B.W. Neuman, D. Penzar, S. Perlman, L.L.M. Poon, D.V. Samborskiy, I.A. Sidorov, I. Sola, J. Ziebuhr, and V. Coronaviridae Study Group of the International Committee on Taxonomy of. 2020. The species Severe acute respiratory syndrome-related coronavirus: classifying 2019-nCoV and naming it SARS-CoV-2. Nature microbiology 5:536–544.

Gustafson, D., S. Raju, R. Wu, C. Ching, S. Veitch, K. Rathnakumar, E. Boudreau, K.L. Howe, and J.E. Fish. 2020. Overcoming Barriers: The Endothelium As a Linchpin of Coronavirus Disease 2019 Pathogenesis? Arterioscler Thromb Vasc Biol 40:1818–1829.

Helms, J., C. Tacquard, F. Severac, I. Leonard-Lorant, M. Ohana, X. Delabranche, H. Merdji, R. Clere-Jehl, M. Schenck, F. Fagot Gandet, S. Fafi-Kremer, V. Castelain, F. Schneider, L. Grunebaum, E. Anglés-Cano, L. Sattler, P.-M. Mertes, F. Meziani, and C.T. Group. 2020. High risk of thrombosis in patients with severe SARS-CoV-2 infection: a multicenter prospective cohort study. Intensive Care Med 46:1089–1098.

Hoffmann, M., H. Kleine-Weber, S. Schroeder, N. Krüger, T. Herrler, S. Erichsen, T.S. Schiergens, G. Herrler, N.H. Wu, A. Nitsche, M.A. Müller, C. Drosten, and S. Pöhlmann. 2020. SARS-CoV-2 Cell Entry Depends on ACE2 and TMPRSS2 and Is Blocked by a Clinically Proven Protease Inhibitor. Cell 181:271–280.e278.

Hu, B., S. Huang, and L. Yin. 2020. The cytokine storm and COVID-19. Journal of Medical Virology n/a:

Huertas, A., D. Montani, L. Savale, J. Pichon, L. Tu, F. Parent, C. Guignabert, and M. Humbert. 2020. Endothelial cell dysfunction: a major player in SARS-CoV-2 infection (COVID-19)? European Respiratory Journal 2001634.

Kumar, S., R. Nyodu, V.K. Maurya, and S.K. Saxena. 2020. Host Immune Response and Immunobiology of Human SARS-CoV-2 Infection. Coronavirus Disease 2019 (COVID-19) 43–53.

Maurer, M., M.S. Gresnigt, A. Last, T. Wollny, F. Berlinghof, R. Pospich, Z. Cseresnyes, A. Medyukhina, K. Graf, M. Gröger, M. Raasch, F. Siwczak, S. Nietzsche, I.D. Jacobsen, M.T. Figge, B. Hube, O. Huber, and A.S. Mosig. 2019. A three-dimensional immunocompetent intestine-on-chip model as in vitro platform for functional and microbial interaction studies. Biomaterials 220:119396.

Ogando, N.S., T.J. Dalebout, J.C. Zevenhoven-Dobbe, R.W. Limpens, Y. van der Meer, L. Caly, J. Druce, J.J.C. de Vries, M. Kikkert, M. Bárcena, I. Sidorov, and E.J. Snijder. 2020. SARS-coronavirus-2 replication in Vero E6 cells: replication kinetics, rapid adaptation and cytopathology. bioRxiv 2020.2004.2020.049924.

Pelaia, C., C. Tinello, A. Vatrella, G. De Sarro, and G. Pelaia. 2020. Lung under attack by COVID-19-induced cytokine storm: pathogenic mechanisms and therapeutic implications. Ther Adv Respir Dis 14:1753466620933508.

Pons, S., S. Fodil, E. Azoulay, and L. Zafrani. 2020. The vascular endothelium: the cornerstone of organ dysfunction in severe SARS-CoV-2 infection. Crit Care 24:353–353.

Rambaut, A., E.C. Holmes, Á. O’Toole, V. Hill, J.T. McCrone, C. Ruis, L. du Plessis, and O.G. Pybus. 2020. A dynamic nomenclature proposal for SARS-CoV-2 lineages to assist genomic epidemiology. Nature microbiology

Ren, Y., T. Shu, D. Wu, J. Mu, C. Wang, M. Huang, Y. Han, X.-Y. Zhang, W. Zhou, Y. Qiu, and X. Zhou. 2020. The ORF3a protein of SARS-CoV-2 induces apoptosis in cells. Cellular & Molecular Immunology

Rennert, K., S. Steinborn, M. Gröger, B. Ungerböck, A.M. Jank, J. Ehgartner, S. Nietzsche, J. Dinger, M. Kiehntopf, H. Funke, F.T. Peters, A. Lupp, C. Gärtner, T. Mayr, M. Bauer, O. Huber, and A.S. Mosig. 2015. A microfluidically perfused three dimensional human liver model. Biomaterials 71:119–131.

Shang, J., Y. Wan, C. Luo, G. Ye, Q. Geng, A. Auerbach, and F. Li. 2020. Cell entry mechanisms of SARS-CoV-2. Proceedings of the National Academy of Sciences 117:11727.

Stertz, S., M. Reichelt, M. Spiegel, T. Kuri, L. Martínez-Sobrido, A. García-Sastre, F. Weber, and G. Kochs. 2007. The intracellular sites of early replication and budding of SARS-coronavirus. Virology 361:304–315.

Tang, X., C. Wu, X. Li, Y. Song, X. Yao, X. Wu, Y. Duan, H. Zhang, Y. Wang, Z. Qian, J. Cui, and J. Lu. 2020. On the origin and continuing evolution of SARS-CoV-2. National Science Review 7:1012–1023.

Thiel, V., and F. Weber. 2008. Interferon and cytokine responses to SARS-coronavirus infection. Cytokine Growth Factor Rev 19:121–132.

Thomas, L., Z. Rao, J. Gerstmeier, M. Raasch, C. Weinigel, S. Rummler, D. Menche, R. Muller, C. Pergola, A. Mosig, and O. Werz. 2017. Selective upregulation of TNFalpha expression in classically-activated human monocyte-derived macrophages (M1) through pharmacological interference with V-ATPase. Biochemical pharmacology 130:71–82.

Thoms, M., R. Buschauer, M. Ameismeier, L. Koepke, T. Denk, M. Hirschenberger, H. Kratzat, M. Hayn, T. Mackens-Kiani, J. Cheng, J.H. Straub, C.M. Stürzel, T. Fröhlich, O. Berninghausen, T. Becker, F. Kirchhoff, K.M.J. Sparrer, and R. Beckmann. 2020. Structural basis for translational shutdown and immune evasion by the Nsp1 protein of SARS-CoV-2. Science (New York, N.Y.) eabc8665.

Varga, Z., A.J. Flammer, P. Steiger, M. Haberecker, R. Andermatt, A.S. Zinkernagel, M.R. Mehra, R.A. Schuepbach, F. Ruschitzka, and H. Moch. 2020. Endothelial cell infection and endotheliitis in COVID-19. The Lancet 395:1417–1418.

Wichmann, D., J.-P. Sperhake, M. Lütgehetmann, S. Steurer, C. Edler, A. Heinemann, F. Heinrich, H. Mushumba, I. Kniep, A.S. Schröder, C. Burdelski, G. de Heer, A. Nierhaus, D. Frings, S. Pfefferle, H. Becker, H. Bredereke-Wiedling, A. de Weerth, H.-R. Paschen, S. Sheikhzadeh-Eggers, A. Stang, S. Schmiedel, C. Bokemeyer, M.M. Addo, M. Aepfelbacher, K. Püschel, and S. Kluge. 2020. Autopsy Findings and Venous Thromboembolism in Patients With COVID-19. Annals of Internal Medicine

Yip, M.S., N.H.L. Leung, C.Y. Cheung, P.H. Li, H.H.Y. Lee, M. Daëron, J.S.M. Peiris, R. Bruzzone, and M. Jaume. 2014. Antibody-dependent infection of human macrophages by severe acute respiratory syndrome coronavirus. Virol J 11:82–82.

